# Gas-sensing neurons prime mitochondrial fitness to offset metabolic stress

**DOI:** 10.1101/2025.06.18.660458

**Authors:** Rebecca Cornell, Ava Handley, Roger Pocock

## Abstract

Animals integrate environmental and internal cues to maintain homeostasis and health. The mitochondrial stress response is an essential cytoprotective mechanism, and priming its activation provides a survival advantage. Here, we show that the *Caenorhabditis elegans* receptor guanylyl cyclase GCY-9 regulates neuropeptide signalling from carbon dioxide sensing neurons to govern a non-canonical mitochondrial stress response in the intestine. This stress response induces atypical mitochondrial chaperone transcription, confers mitochondrial stress resistance, and increases mitochondrial membrane potential and respiration. GCY-9 loss disrupts pathogen avoidance, leading to indiscriminate feeding. We show that starvation decreases GCY-9 expression and propose that the resultant cytoprotective program is launched to offset risks associated with this behaviour. Thus, environmental sensing by peripheral neurons can pre-emptively enhance systemic mitochondrial function in response to metabolic uncertainty.

**One-Sentence Summary:** Protecting mitochondria by integrating environmental signals

## Main Text

Coordinating stress responses across tissues is essential for cellular homeostasis and organismal health. Cells employ sophisticated mechanisms to detect and respond to stressors, including the organelle specific mitochondrial and endoplasmic reticulum unfolded protein responses (UPR^mt^ and UPR^ER^) and the cytosolic heat shock response. The ability to anticipate stressors and prime stress responses would offer a powerful cellular and organismal survival advantage. The nervous system is critical for sensing environmental challenges and coordinating stress responses in distal tissues (*1-5*). Anticipatory signalling offers clear adaptive benefits, especially under fluctuating environmental conditions, however, we understand little of the neuronal and molecular mechanisms that orchestrate this cross-talk.

The *Caenorhabditis elegans* gas sensing BAG neurons are optimally positioned at the intersection of environmental sensing and physiological adaptability. The BAG neurons are activated by elevated carbon dioxide (CO_2_) levels (*6*), which can influence acid-base homeostasis, cellular respiration, and oxidative stress – all of which are associated with mitochondrial health. Furthermore, our previous studies showed that the BAG neurons influence metabolism and animal behaviour through the ETS-5 transcription factor, an orthologue of mammalian FEV/Pet1 (*7, 8*). Due to their ability to integrate external and internal environmental cues, we posited the BAG neurons as candidates for anticipating and orchestrating anticipatory mitochondrial stress responses.

### Loss of the neuronal transcription factor ETS-5 from the BAG neurons induces a non-canonical UPR^mt^

The terminal fate of the BAG neurons – particularly relating to their roles in environmental sensing and systemic metabolism – is mediated by the ETS-5 transcription factor (*9, 10*). We analysed the *hsp-6p::gfp* mitochondrial chaperone reporter to determine whether ETS-5 influences the systemic UPR^mt^. HSP-6/mtHsp70 is required for importing nuclear-encoded proteins into the mitochondrial matrix and is a key target of the UPR^mt^. We found that two independent *ets-5* deletion mutants exhibited increased intestinal *hsp-6p::gfp* expression (Fig. 1A and B). Typically, UPR^mt^ proteins are upregulated in concert (*11-13*). However, we found that a second UPR^mt^ reporter for the mitochondrial matrix chaperonin HSP-60 (*hsp-60p::gfp)*, showed decreased intestinal expression in the *ets-5* deletion mutants (Fig. 1D and E). Aberrant *hsp-6p::gfp* and *hsp-60p::gfp* expression in *ets-5* mutant animals was rescued by driving *ets-5* cDNA with the BAG neuron specific *gcy-9* promoter (Fig. 1C and F). Canonically, *hsp-6* and *hsp-60* transcription is upregulated in the UPR^mt^ by the master transcription factor ATFS-1/ATF5, supplemented by the homeodomain transcription factor DVE-1/SATB and its co-activator UBL-5/Hub1 (*11, 14, 15*). To examine the requirement of these factors for *hsp-6p::gfp* upregulation in *ets-5* mutant animals, we performed RNA-mediated interference (RNAi) knockdown of ATFS-1, DVE-1 and UBL-5. We used an *unc-25* mutant as a positive control for canonical *hsp-6p::gfp* induction (*5*). We found that *atfs-1* and *ubl-5*, but not *dve-1*, are required for the *hsp-6p::gfp* induction observed in *ets-5* mutant animals (Fig. 1G). To confirm DVE-1 independence, we analysed a DVE-1 fusion protein (DVE-1::GFP) which localises to nuclei upon activation (Fig. S1) (*15*). We found no difference in the number of intestinal nuclear puncta in unstressed or stressed conditions (Fig 1H and Fig S1A) or in the DVE-1 nuclear:cytosolic fluorescence ratio (Fig. S1B-C) between wild-type and *ets-5* mutant animals. These data imply that DVE-1 is not required for the atypical mitochondrial stress response observed in *ets-5* mutant animals.

**Fig. 1.**
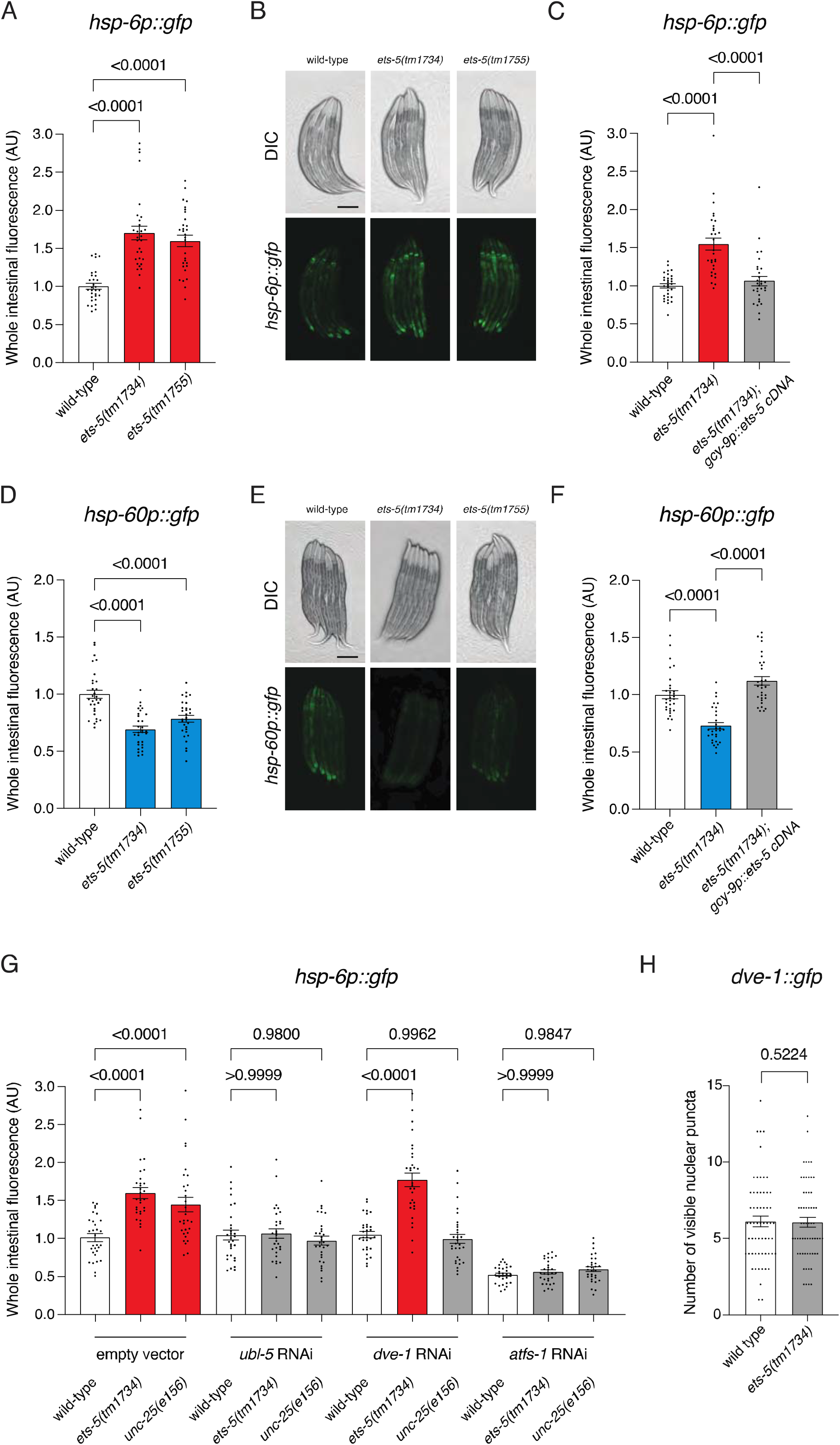
Loss of the neuronal transcription factor ETS-5 from the BAG neurons induces a non-canonical UPR^mt^. **A, B** Quantification (**A**) and DIC/fluorescent micrographs (**B**) of UPR^mt^ reporter (*hsp-6p::gfp*) expression in L4 larvae of wild-type, *ets-5(tm1734)* and *ets-5(tm1755)* animals. (**C)** Quantification of UPR^mt^ reporter (*hsp-6p::gfp*) expression in L4 larvae of wild-type, *ets-5(tm1734)*, and *ets-5(tm1734); gcy-9p::ets-5 cDNA* animals. **D, E** Quantification (**D**) and DIC/fluorescent micrographs (**E**) of UPR^mt^ reporter (*hsp-60p::gfp*) expression in L4 larvae of wild-type, *ets-5(tm1734)* and *ets-5(tm1755)* animals. (**F)** Quantification of UPR^mt^ reporter (*hsp-60p::gfp*) expression in L4 larvae of wild-type, *ets-5(tm1734)*, and *ets-5(tm1734); gcy-9p::ets-5 cDNA* animals. (**G**) Quantification of UPR^mt^ reporter (*hsp-6p::gfp*) expression in L4 larvae of wild-type and *ets-5(tm1734)* animals grown on empty vector (EV), *ubl-5, dve-1* or *atfs-1* RNAi from hatch. (**H**) Quantification of DVE-1::GFP nuclear puncta in wild-type and *ets-5(tm1734)* animals. *n*□=□30 animals (**A, C, D, F** and **G**) or *n* = 60 animals (**H**). *P* values assessed by one-way analysis of variance (ANOVA) with Tukey’s post hoc test (**A, C, D, F** and **G**), or unpaired t test with Welch’s correction (**H**). Error bars indicate SEM. Scale bar, 250□µm.

Cellular stress responses can be independently activated or integrated across multiple organelles and cellular compartments. We therefore analysed the ER stress response marker *hsp-4p::gfp* (HSPA6/BiP), the cytosolic stress marker *hsp-16*.*2p::gfp* (HSBP1), and two oxidative stress markers *sod-3p::gfp* (SOD2) and *gst-4p::gfp* (GSTA4) in *ets-5* mutant animals. We detected no change in ER stress (Fig. S2A and B), cytosolic stress (Fig. S2C and D) or oxidative stress (Fig. S2E-H) responses in *ets-5* mutants. Together, these data reveal that the BAG neurons control a highly specific and non-canonical systemic UPR^mt^ (HSP-6 upregulation and HSP-60 downregulation) through the ETS-5 transcription factor.

### Loss of ETS-5 improves mitochondrial health readouts

As HSP-6/HSP-60 expression in *ets-5* mutant animals is atypical, we aimed to determine how mitochondrial fitness is impacted. First, we performed an acute paraquat survival assay to assess mitochondrial stress resistance in *ets-5* mutant animals. Paraquat is a herbicide that disrupts complex I of the electron transport chain, resulting in increased superoxide levels and organismal death (*16*). We found that *ets-5* mutant animals were resistant to paraquat exposure compared to wild-type animals (Fig. 2A). Wild-type levels of paraquat resistance were restored to *ets-5* mutant animals by resupplying *ets-5* cDNA specifically to the BAG neurons (Fig. 2A). Second, we assessed mitochondrial membrane potential in *ets-5* mutant animals using tetramethylrhodamine ethyl ester (TMRE) staining. Higher membrane potential, which is beneficial for mitochondrial function, leads to higher TMRE absorption into the mitochondrial matrix (*17*). We found that *ets-5* mutant animals exhibit increased TMRE fluorescence intensity (Fig. 2B and C), suggesting increased mitochondrial membrane potential. Wild-type levels of TMRE fluorescence intensity were restored to *ets-5* mutant animals by resupplying *ets-5* cDNA specifically to the BAG neurons (Fig. 2B and C). Finally, we found that loss of *ets-5* increases both basal and maximal oxygen (O_2_) consumption rates (Fig. 2D-F), suggesting increased capacity for energy production. Together, these data indicate that the atypical stress response induced by ETS-5 loss from the BAG neurons is beneficial for systemic mitochondrial fitness, both at rest and under stress.

**Fig. 2.**
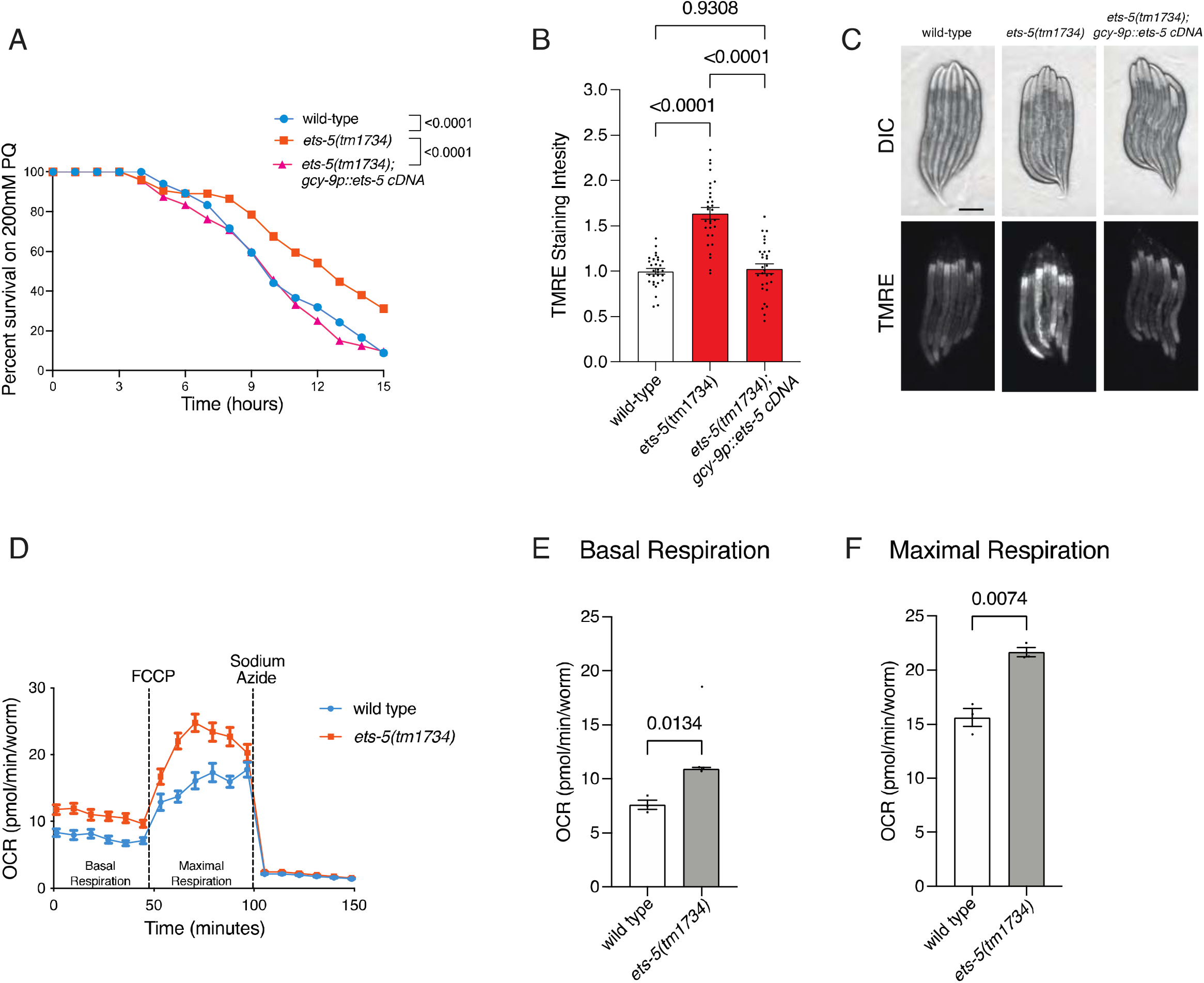
Loss of ETS-5 improves mitochondrial health readouts. (**A**) Survival analysis of wild-type, *ets-5(tm1734)*, and *ets-5(tm1734); gcy-9p::ets-5 cDNA* animals exposed to 200□mM paraquat from the L4 larval stage. *n*□=□66, 74, 72 animals (top to bottom). **B, C** Quantification (**B**) and DIC/fluorescent micrographs (**C**) of mitochondrial membrane potential reporter TMRE staining intensity in L4 larvae of wild-type, *ets-5(tm1734)*, and *ets-5(tm1734); gcy-9p::ets-5 cDNA* animals. *n* = 30. Scale bars, 250μm. **D-F** Seahorse oxygen consumption rate (OCR) assay. (**D**) Kinetic OCR analysis of wild-type and *ets-5(tm1734)* animals, measuring (**E**) basal, and (**F**) maximal oxygen consumption. *n* = 3 replicates (∼100 animals per replicate). *P* values assessed by two-way analysis of variance (ANOVA) with Tukey’s post hoc test (**A**) one-way ANOVA with Tukey’s post hoc test (**B**), or unpaired t test with Welch’s correction (**E** and **F**). Error bars indicate SEM.

### BAG-derived neuropeptides mediate intestinal mitochondrial stress responses

How does ETS-5, acting in the BAG neurons, impact mitochondrial health in the non-innervated *C. elegans* intestine? We hypothesized that specific neuropeptides may act as BAG-intestinal signals in this context. To investigate this, we used the auxin inducible degron system to degrade UNC-31/CAPS – a phosphoinositide-binding protein required for dense core vesicle release – specifically in the BAG neurons by expressing TIR1 under the BAG specific *gcy-9* promoter, as we performed previously (*18*). When these animals are fed auxin, neuropeptide release is inhibited specifically from the BAG neurons. We found that auxin exposure increased *hsp-6p::gfp* and decreased *hsp-60p::gfp* expression – phenocopying *ets-5* mutant animals (Fig. 3A and B). This implies that the BAG neurons utilise neuropeptides to control systemic mitochondrial health.

**Fig. 3.**
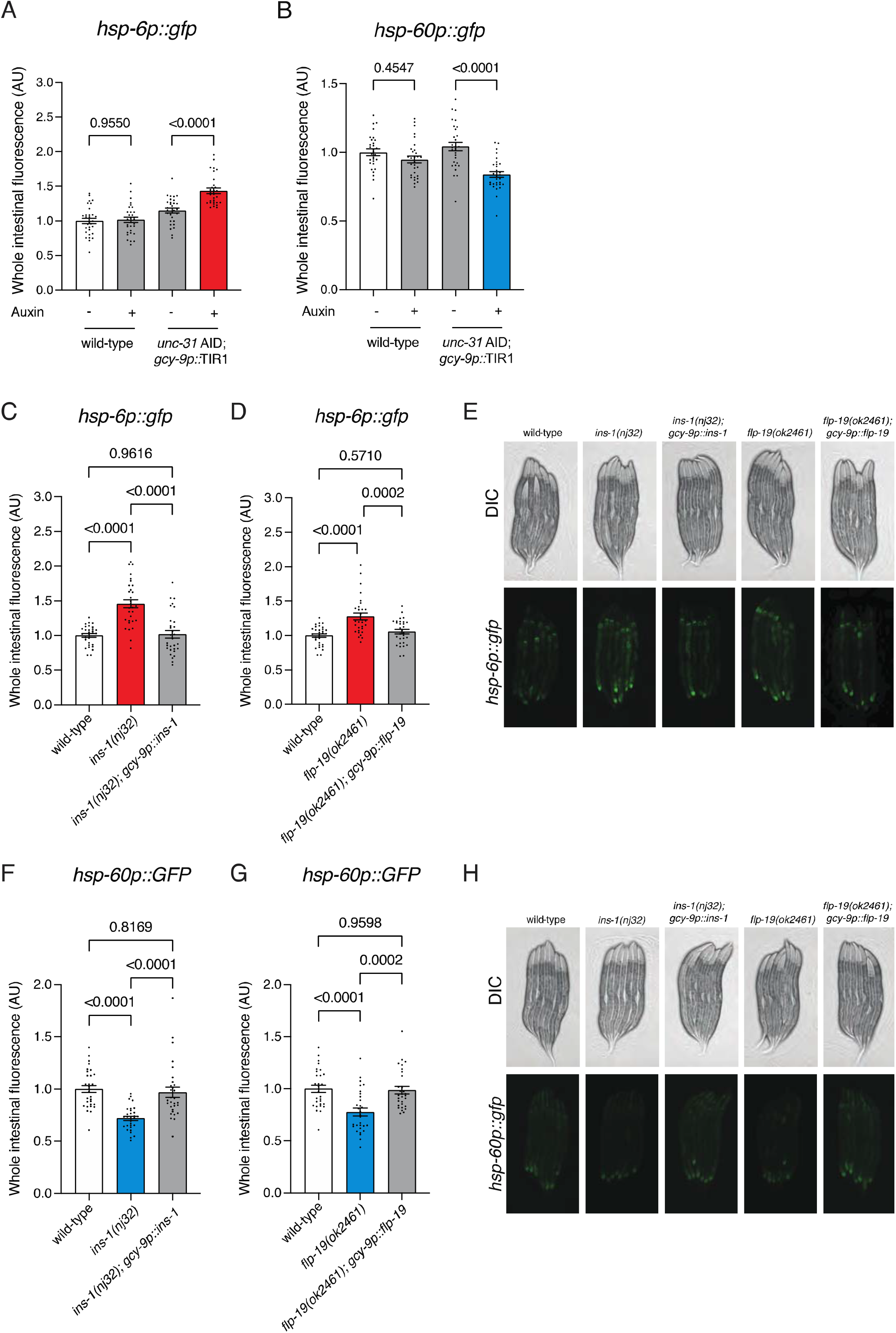
BAG-derived neuropeptides mediate intestinal mitochondrial stress responses. (**A, B**) Quantification of UPR^mt^ reporter (*hsp-6p::gfp*) (**A**) and (*hsp-60p::gfp*) (**B**) expression in L4 larvae of wild-type and *unc-31 AID; gcy-9p::TIR1* animals, with and without auxin treatment. **C-E** Quantification (**C** and **D**) and DIC/fluorescent micrographs (**E**) of UPR^mt^ reporter (*hsp-6p::gfp*) expression in L4 larvae of wild-type, *ins-1(nj32)* and *gcy-9p::ins-1; ins-1(nj32)* (**C**) and wild-type, *flp-19(ok2461)* and *gcy-9p::flp-19; flp-19(ok2461)* (**D**) animals. **F-H** Quantification (**F** and **G**) and DIC/fluorescent micrographs (**H**) of UPR^mt^ reporter (*hsp-60p::gfp*) expression in L4 larvae of wild-type, *ins-1(nj-32)* and *gcy-9p::ins-1; ins-1(nj32)* (**F**) and wild-type, *flp-19(ok2461)* and *gcy-9p::flp-19; flp-19(ok2461)* (**G**) animals. *n* = 30. *P* values assessed by one-way analysis of variance (ANOVA) with Tukey’s post hoc test. Error bars indicate SEM. Scale bar, 250□µm.

To determine which BAG-expressed neuropeptides control the distal mitochondrial stress response, we screened neuropeptide mutants for changes to *hsp-6p::gfp* and *hsp-60p::gfp* expression (Fig. S3). We screened insulin-like peptides (INS-1, INS-14 and INS-29) and FMRF-like peptides (FLP-4, FLP-6, FLP-10, FLP-12, FLP-13, FLP-17 and FLP-19), all of which (except FLP-17) are expressed in multiple neurons in addition to the BAG neurons. Of these genes, loss of *flp-4* was the only neuropeptide mutant that showed a significant increase in both *hsp-6p::gfp* and *hsp-60p::gfp* levels (Fig. S3C and D). Interestingly, *ins-1* and *flp-19* mutants both showed increased *hsp-6p::gfp* and decreased *hsp-60p::gfp* expression. As both *ins-1* and *flp-19* are transcriptionally regulated by ETS-5 in the BAG neurons (*7, 8*), and their combined loss causes UPR^mt^ dysregulation that is equal to loss of *ets-5* (Fig. S4), we proposed that the BAG neurons utilise INS-1 and FLP-19 to mediate systemic mitochondrial function. To confirm this, we resupplied a single copy of *ins-1* and *flp-19* genomic DNA specifically to the BAG neurons in the respective mutants, and found that this rescued the increased *hsp-6p::gfp* (Fig. 3C-E) and decreased *hsp-60p::gfp* (Fig. 3F-H) expression.

### ETS-5 utilises BAG-derived INS-1 and FLP-19 to mediate systemic mitochondrial health

As *ins-1* and *flp-19* mutant animals exhibit the same atypical chaperone protein expression pattern as *ets-5* mutants, we examined whether this translated to the same enhanced mitochondrial fitness. We measured TMRE staining intensity and found that loss of both *ins-1* and *flp-19* was equivalent, but not additive, to loss of *ets-5* in increasing mitochondrial membrane potential (Fig. 4A-B). We also found that loss of *ins-1* or *flp-19* conferred paraquat resistance (Fig. S5A and B), and the *ins-1; flp-19* double mutant was equivalent to loss of *ets-5* (Fig. 4C). Conversely, loss of *flp-4*, which led to increased *hsp-6p::gfp* and *hsp-60p::gfp* expression, did not affect paraquat sensitivity (Fig. S5C). Whereas, loss of *flp-17*, which caused increased *hsp-6p::gfp*, and no change to *hsp-60p::gfp*, resulted in a mild resistance to paraquat (Fig. S5D).

**Fig. 4.**
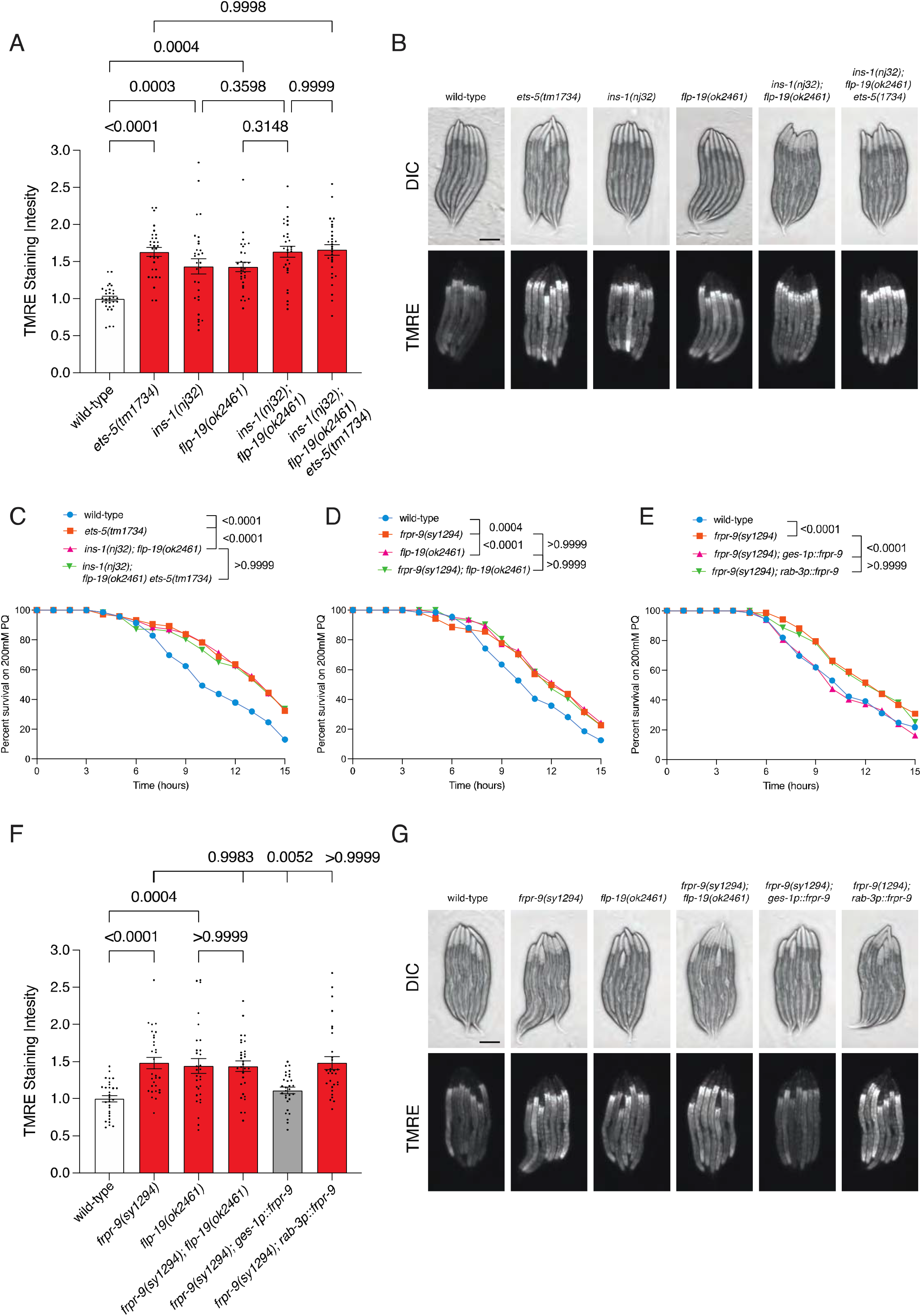
ETS-5 utilises BAG-derived INS-1 and FLP-19 to mediate systemic mitochondrial health. **A, B** Quantification (**A**) and DIC/fluorescent micrographs (**B**) of mitochondrial membrane potential reporter TMRE staining intensity in L4 larvae of wild-type, *ets-5(tm1734), ins-1(nj32); flp-19(ok2461), ins-1(nj32); flp-19(ok2461)* and *ins-1(nj32); flp-19(ok2461) ets-5(tm1734)* animals. *n* = 30. Scale bars, 250μm. (**C**) Survival analysis of wild-type, *ets-5(tm1734), ins-1(nj32); flp-19(ok2461)* and *ins-1(nj32); flp-19(ok2461) ets-5(tm1734)* animals exposed to 200□mM paraquat from the L4 larval stage. n□=□69, 74, 70, 71 animals (top to bottom). (**D**) Survival analysis of wild-type, *frpr-9(sy1294), flp-19(ok2461)* and *frpr-9(sy1294); flp-19(ok2461)* animals exposed to 200□mM paraquat from the L4 larval stage. n□= 65, 67, 66, 72 animals (top to bottom). (**E**) Survival analysis of wild-type, *frpr-9(sy1294), ges-1p::frpr-9; frpr-9(sy1294)* and *rab-3p::frpr-9; frpr-9(sy1294)* animals exposed to 200□mM paraquat from the L4 larval stage. *n*□=□65, 68, 67, 68 animals (top to bottom). **F, G** Quantification (**F**) and DIC/fluorescent micrographs (**G**) of mitochondrial membrane potential reporter TMRE staining intensity in L4 larvae of wild-type, *frpr-9(sy1294), flp-19(ok2461), frpr-9(sy1294)*; *flp-19(ok2461), ges-1p::frpr-9; frpr-9(sy1294)* and *rab-3p::frpr-9; frpr-9(sy1294)* animals. *n* = 30. *P* values assessed by two-way analysis of variance (ANOVA) with Tukey’s post hoc test (**A, D** and **E**) or one-way ANOVA with Tukey’s post hoc test (**B** and **F**). Error bars indicate SEM. Scale bars, 250μm.

### BAG derived FLP-19 meditates systemic mitochondrial health via intestinal receptor FRPR-9

DAF-2 is the only insulin-like growth factor receptor in *C. elegans*, with well-documented roles relating to mitochondrial function that stem from many pathways. For example, DAF-2 inactivation in the intestine promotes paraquat resistance (*19*). We therefore focused our research on the recently discovered FLP-19 receptor, FRPR-9 (*20*). We found that *frpr-9* loss induces increased mitochondrial stress resistance (Fig. 4D) and increased mitochondrial membrane potential (Fig. 4F-G). Importantly, these phenotypes were not additive to loss of *flp-19* (Fig. 4D, F-G). FRPR-9 is expressed in both neurons and intestinal cells (*21*). To determine where FRPR-9 acts to mediate intestinal mitochondrial health, we resupplied *frpr-9* genomic DNA in the intestine (*ges-1* promoter) or pan-neuronally (*rab-3* promoter). We found that intestinal *frpr-9* expression, but not neuronal *frpr-9* expression, restored wild-type paraquat survival (Fig. 4E) and mitochondrial membrane potential to *frpr-9* mutant animals (Fig. 4F-G). These data imply that FLP-19 released from the BAG neurons binds to FRPR-9 on intestinal cells to mediate systemic mitochondrial health.

### BAG carbon dioxide machinery controls systemic mitochondrial health

We previously showed that *flp-19* expression in the BAG neurons depends on the CO□-sensing receptor guanylate cyclase GCY-9, a transcriptional target of ETS-5 (*22*). We therefore wondered if *ins-1* expression is also influenced by *gcy-9* loss. We found that the number of BAG neurons expressing an *ins-1* transcriptional reporter is drastically reduced in a *gcy-9* deletion mutant (Fig. 5A and B). We therefore hypothesised that GCY-9 – and therefore the ability to sense CO_2_ – may be mechanistically involved in the BAG-mediated mitochondrial stress response. Indeed, we found that *gcy-9* loss caused increased *hsp-6p::gfp*, and decreased *hsp-60p::gfp* expression, which was rescued by expressing *gcy-9* cDNA specifically in the BAG neurons (Fig. 5C and D). A *gcy-9 ets-5* double mutant was not additive to either single mutant with respect to *hsp-6p::gfp* and *hsp-60p::gfp* regulation (Fig. S6), indicating that *ets-5* and *gcy-9* act in the same pathway in the BAG neurons to mediate the systemic UPR^mt^. Furthermore, *gcy-9* loss conferred increased mitochondrial stress resistance (Fig. 5E) and increased mitochondrial membrane potential, phenotypes that were not enhanced by simultaneous *ets-5* loss (Fig. 5F and G). Intriguingly, and consistent with previous studies, we found that *gcy-9* loss did not increase intestinal fat storage (*23*) (Fig. S7), in contrast to *ets-5* mutants, which accumulate excess intestinal fat (*7*). Furthermore, *ets-5* expression is not influenced by loss of *gcy-9* (Fig. S8). These results imply that increased fat storage and the associated behavioural phenotypes observed in *ets-5* mutant animals exist in a separate, but potentially linked, molecular pathway to the stress response pathway described here.

**Fig. 5.**
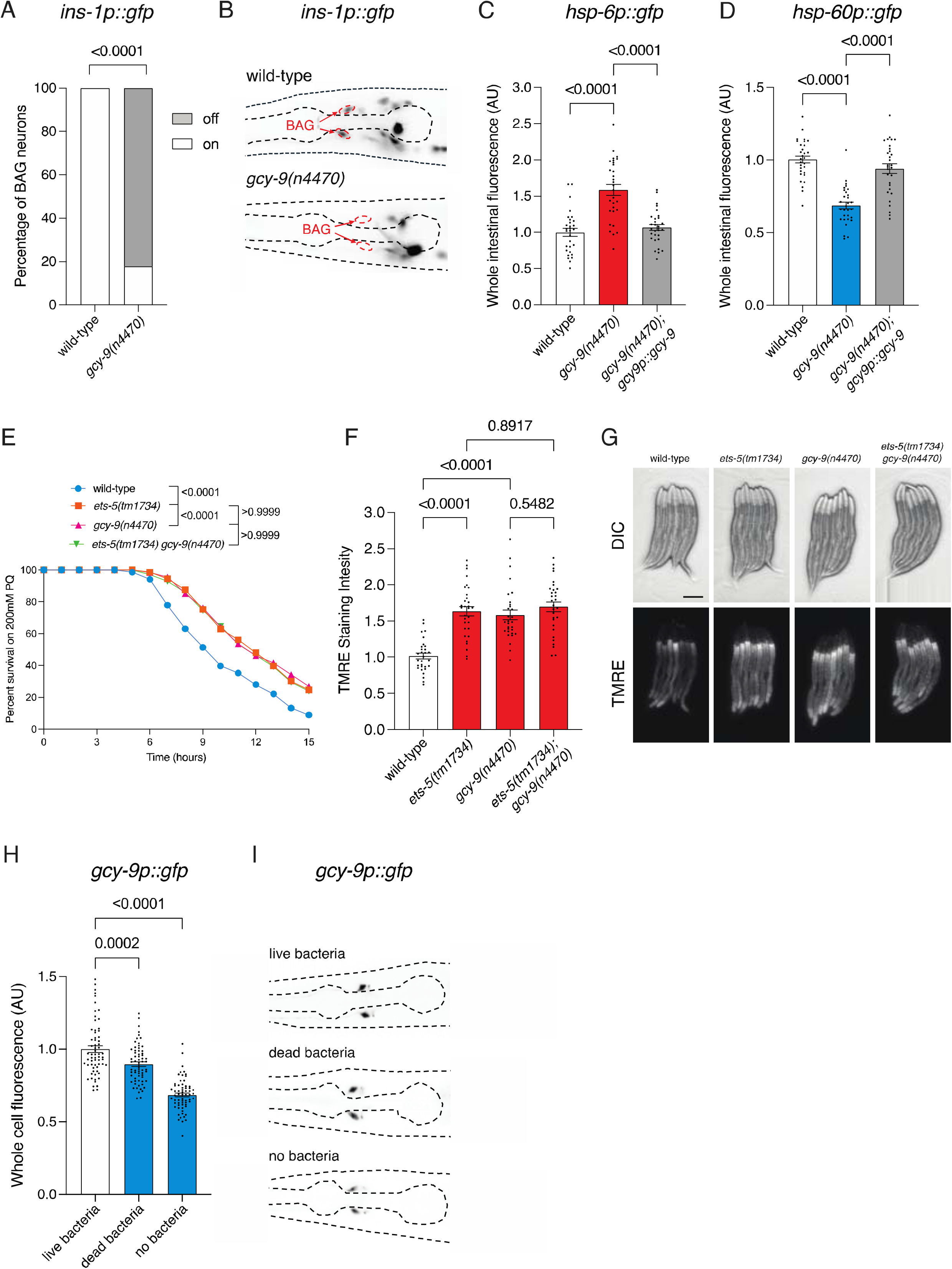
BAG carbon dioxide machinery controls systemic mitochondrial health. **A, B** Quantification (**A**) and fluorescent micrographs (**B**) of *ins-1p::gfp* reporter expression in the BAG neurons in L4 larvae of wild-type and *gcy-9(n4470)* animals. *n* = 30. Scale bar, 50μm. (**C**) Quantification of UPR^mt^ reporter (*hsp-6p::gfp*) expression in L4 larvae of wild-type, *gcy-9(n4470)* and *gcy-9p::gcy-9; gcy-9(n4470)* animals. *n* = 30. (**D**) Quantification of UPR^mt^ reporter (*hsp-60p::gfp*) expression in L4 larvae of wild-type, *gcy-9(n4470)* and *gcy-9p::gcy-9; gcy-9(n4470)* animals. *n* = 30. (**E**) Survival analysis of wild-type, *ets-5(tm1734), gcy-9(n4470)* and *ets-5(tm1734) gcy-9(n4470)* animals exposed to 200□mM paraquat from the L4 larval stage. *n*□= 68, 73, 67, 71 animals (top to bottom). **F, G** Quantification (**F**) and DIC/fluorescent micrographs (**G**) of mitochondrial membrane potential reporter TMRE staining intensity in L4 larvae of wild-type, *ets-5(tm1734), gcy-9(n4470)* and *ets-5(tm1734) gcy-9(n4470)* animals. *n* = 30. Scale bar, 250μm. **H, I** Quantification (**H**) and DIC/fluorescent micrographs (**I**) of *gcy-9p::gfp* reporter expression in BAG neurons in L4 larvae of wild-type exposed to live, dead or no bacteria for 24 hours. *n* = 30. Scale bar, 50μm. *P* values assessed by unpaired t test with Welch’s correction (**A**) one-way ANOVA with Tukey’s post hoc test (**B**), or (**C, D, F** and **H**) or two-way analysis of variance (ANOVA) with Tukey’s post hoc test (**E**). Error bars indicate SEM.

CO_2_ is a metabolic by-product, and fluctuations in environmental CO_2_ can signal key environmental factors such as food quality, overcrowding, and hypoxia. Why would impaired CO_2_ sensing enhance stress resistance? We propose that loss of peripheral CO_2_ sensing functions as a “blackout alarm”, reflecting a failure to detect environmental status and triggering a primed stress response in high metabolic tissues such as the intestine. The unique stress response described here likely represents an adaptive failsafe triggered to ensure survival in ephemeral environments.

Previous studies showed that *gcy-9* mutants fail to avoid pathogenic bacteria (*24*). Combined with our data, this raises the possibility that *gcy-9* expression – under the transcriptional control of ETS-5 – is dynamically regulated in response to environmental cues such as food. We thus wondered if, during starvation, animals might downregulate *gcy-9* expression in the BAG neurons to suppress avoidance of potentially pathogenic but nutritionally valuable bacteria. Indeed, we found that *gcy-9* transcriptional reporter expression decreased following starvation or exposure to dead, metabolically inactive bacteria (Fig. 5H and I). We propose that this transcriptional mechanism balances feeding with cellular protection and exists to buffer the risk associated with non-discriminatory feeding when food is scarce. By suppressing *gcy-9* expression, animals increase their likelihood of consuming available food, while simultaneously activating an atypical systemic mitochondrial stress response that primes tissues to offset the potential harm of metabolic stress. This hypothesis is supported by previous research, revealing that FRPR-9 loss enhanced resistance to both gram negative and gram-positive pathogenic bacteria (*25*).

## Discussion

Here, we identify a pair of peripheral sensory neurons (the *C. elegans* gas-sensing BAG neurons) that mediate protective plasticity through non-autonomous priming of the intestinal mitochondrial stress response, enabling organismal resistance to metabolic stress. Within the BAG neurons, the ETS-5–GCY-9 regulatory axis controls expression of the INS-1 and FLP-19 neuropeptides that signal to the intestine. We show that the FLP-19 receptor FRPR-9 acts in the intestine to modulate mitochondrial membrane potential and stress resistance.

The specificity of the ETS-5-controlled UPR^mt^ may be due to a distinctive combination of transcription regulators used that does not involve the DVE-1 homeodomain transcription factor. DVE-1 and UBL-5 function together to positively regulate both *hsp-6* and *hsp-60* transcription (*15*). However, previous studies have shown that DVE-1 can act independently of UBL-5 to induce the UPR^mt^ and regulate lifespan (*26, 27*). UBL-5 can also act independently of DVE-1 in yeast (*28*), recently confirmed in *C. elegans* (*29*), where it functions in pre-mRNA alternative splicing. This mechanism may be conserved in humans, as human UBL5 interacts with the cyclin-like kinase CLK4, which plays an important role in spliceosome formation (*30*). These studies provide evidence to support additional roles for UBL-5 outside of DVE-1 coactivation in *C. elegans*.

In our model, we propose that loss of CO□ detection through GCY-9 in the BAG neurons functions as a “blackout alarm,” prompting anticipatory stress responses in metabolically active tissues such as the intestine. Priming stress responses to sensory deficits has been observed in other systems. In mice, ablation of olfactory sensory neurons promotes resistance to diet induced obesity and improves insulin sensitivity, suggesting that sensory input impacts systemic metabolism (*31*).

Integration of environmental and metabolic cues by the BAG neurons to mediate systemic physiology suggests an intriguing evolutionary parallel to mammalian carotid bodies. The carotid bodies are chemoreceptors located bilaterally at the bifurcation of the common carotid arteries that detect changes in arterial blood CO_2_ and O_2_. They also respond to metabolic signals such as blood glucose, insulin, and adrenalin (*32-34*). Overactivation of the carotid bodies during sleep apnoea is associated with increased fasting glucose, insulin resistance, reduced glucose tolerance and impaired pancreatic β cell function (*32, 35, 36*). Carotid body ablation restores insulin sensitivity and glucose tolerance in rats with diet-induced metabolic syndrome (*37*), and denervation prevents the development of hypertension and insulin resistance induced by hypercaloric diets (*33*). However, bilateral carotid body resection confers a significant risk of O_2_ desaturation during mild hypoxia (*38*). Therefore, research into pharmacological modification of key biomolecular pathways in peripheral chemosensors controlling systemic metabolism – such as described here – is required. The BAG neurons also share functional similarities with pancreatic β cells. ETS-5 directly regulates transcription of the insulin-like peptide gene, *ins-1*, in the BAG neurons (*8*). Likewise, the mammalian ETS-5 orthologue – Pet1 – positively regulates insulin expression in mouse pancreatic β cells (*39*). Thus, appropriate regulation of insulin signalling is likely essential for maintaining systemic mitochondrial and metabolic function across evolution.

These functional comparisons to the carotid bodies and pancreatic β cells position the BAG neurons as a powerful evolutionary model for investigating how sensory perception, behaviour and systemic mitochondrial fitness are integrated. Collectively, our findings underscore the nervous system’s capacity to prime protective mitochondrial responses in peripheral tissues by interpreting changes in sensory information.

## Acknowledgments

We thank members of the Pocock laboratory for advice and comments on the manuscript. Imaging for this project was performed at Monash Microimaging. Some strains were provided by the *Caenorhabditis* Genetics Center (University of Minnesota), which is funded by the NIH Office of Research Infrastructure Programs (P40 OD010440) and National BioResource Project of Japan.

## Author contributions

Conceptualization: RC, AH, RP

Methodology: RC, AH, RP

Investigation: RC, AH, RP

Visualization: RC, AH, RP

Funding acquisition: RP, AH

Project administration: RP

Supervision: AH, RP

Writing – original draft: RC

Writing – review & editing: AH, RP

## Data and materials availability

All data is available in the main text or supplementary materials. In addition, Source Data are provided with this paper. There are no accession codes, unique identifiers, or weblinks in our study and no restrictions on data availability. Materials will be available upon request from the Pocock laboratory.

## Supplementary Materials

Materials and Methods Figs. S1 to S8

Table S1

References (*18, 40-47*) Data S1

