## Supplementary Information for "Gas-sensing neurons prime mitochondrial fitness to offset metabolic stress"

### Materials and Methods

#### *C. elegans* culture

All *C. elegans* strains were grown on nematode growth media (NGM) plates seeded with 400µL OP50 *Escherichia coli.* Strains were backcrossed to N2 a minimum of three times and maintained well fed at a constant temperature of 20°C for a minimum of two generations prior to analysis. Strains used in this study are listed in Table S1.

#### Fluorescence microscopy and quantification

Worms were anesthetised in 0.1ng/ml levamisole on a 5% agarose pad. Images for analysis were taken using a Zeiss AXIO Imager M2 upright fluorescence microscope and ZEN 2.0 software. Image J software was used to determine whole intestinal / whole cell fluorescence by isolating the intestine or BAG neuron area and measuring integrated density (IntDensity). The corrected total cellular fluorescence was calculated using the following formula: CTCF = Intestinal IntDensity – (area * background MeanGrey).

Localisation of the DVE-1 fusion protein, DVE-1::GFP, was measured by counting visible nuclear puncta across the intestine, as described previously *(40).* To confirm that analysis of DVE-1 nuclear localisation was possible in the strains analysed, worms were heat shocked for 30 minutes and recovered for 1 hour before scoring. To measure nuclear:cytosolic fluorescence ratio, the nuclei of the first two intestinal cells were traced and fluorescence measured and compared to cytosolic fluorescence.

For the *ins-1* transcriptional reporter, visible fluorescence in the BAG neurons was recorded as "on", and no visible fluorescence in the BAG neurons was recorded as "off".

#### RNAi-mediated interference (RNAi) by feeding

RNAi was performed by feeding as described previously *(41)*. Briefly, HT115 bacteria containing control (L4440) or experimental (containing targeted dsRNA) plasmids was grown overnight at 37°C in LB media. IPTG (Isopropyl β- d-1-thiogalactopyranoside)-containing plates were also dried overnight at 37°C. Plates were then uniformly seeded with overnight cultures and dried at 37°C. Eggs for experimental strains were placed on seeded RNAi plates and grown to L4 at 20°C over 48 hours before being imaged for analysis.

#### *C. elegans* single copy rescue strain generation

Single copy rescue strains were generated using self-excising drug selection cassette (SEC) mediated CRISPR/Cas9, as previously described *(42)*. Plasmids were confirmed by sequencing prior to injection. Microinjection into young adult animals was performed using standard methods.

gcy-9p::ins-1 single copy rescue construct

The 1239bp *ins-1* DNA was amplified from a *C. elegans* genomic DNA library with forward (TGAGATGATAAATCGATATGTACTGGTTTCGTCAAGTTTAC) and reverse (GCATTTATCTTAGCCGGCTCAATTATCGTCCTGATTGCAGC) primers from Sigma-Aldrich incorporating *Bsu15I-PdiI* restriction sites. The *gcy-9p-FLP-p2A-H2B-mTurq2* plasmid was digested using *Bsu15I-PdiI* and the *ins-1* DNA was inserted using the In-Fusion HD Cloning Kit (Takara Bio). The resultant *gcy-9p::ins-1 DNA* plasmid was injected at 10ng/µl, with 50ng/µl pCFJ2474 (Cas9-expression), 50ng/µl pJW1882 (ttTi4348 targeting sequence) and 5ng/µl *myo-2p::mCherry into ins-1(nj32); hsp-6p::gfp* and *ins-1(nj32); hsp-60p::gfp* animals.

gcy-9p::flp-19 single copy rescue construct

The 746bp *flp-19* DNA was amplified from a *C. elegans* genomic DNA library with forward (TGAGATGATAAATCGATATGTCCTTCCAGCTAACGC) and reverse (GCATTTATCTTAGCCGGCTTATCCGAACCTGACTGAGC) primers from Sigma-Aldrich incorporating *Bsu15I-PdiI* restriction sites. The *gcy-9p-FLP-p2A-H2B-mTurq2* plasmid was digested using *Bsu15I-PdiI* and the *flp-19* DNA was inserted using the In-Fusion HD Cloning Kit (Takara Bio). The resultant *gcy-9p::flp-19 DNA* plasmid was injected at 10ng/µl, with 50ng/µl pCFJ2474 (Cas9 expression), 50ng/µl pJW1882 (ttTi4348 targeting sequence) and 5ng/µl *myo-2p::mCherry into flp-19(ok2461); hsp-6p::gfp* and *into flp-19(ok2461); hsp-60p::gfp* animals.

ges-1p::frpr-9 single copy rescue construct

The 2719bp *frpr-9* DNA was amplified from a *C. elegans* genomic DNA library with forward (TTCAGATCGATATCGATATGACATCTACTGTCATCTACG) and reverse (GCATTTATCTTAGCCGGCTTAGGCGAGAGTGGCAAATCG) primers from Sigma-Aldrich incorporating *Bsu15I-PdiI* restriction sites. The *ges-1p-GFP-C1-H2B-SEC* plasmid was digested using *Bsu15I-PdiI* to remove the GFP and the *frpr-9* DNA was inserted using the In-Fusion HD Cloning Kit (Takara Bio). The resultant *ges-1p::frpr-9 DNA* plasmid was injected at 10ng/µl, with 50ng/µl pCFJ2474 (Cas9 expression), 50ng/µl pJW1882 (ttTi4348 targeting sequence) and 5ng/µl *myo-2p::mCherry into frpr-9(sy1294)* animals.

rab-3p::frpr-9 single copy rescue construct

The 4382bp *rab-3* promoter was amplified from a *C. elegans* genomic DNA library with forward (ACGGCCAGTGCGGCCCGAGTTTTGACTGGCTTTCA) and reverse (GTAGATGTCATATCGATCTGAAAATAGGGCTACTGTAGATTT) primers from Sigma-Aldrich incorporating *NotI-Bsu15I* restriction sites. The *ges-1p::frpr-9* plasmid was digested using *NotI-Bsu15I* to remove the *ges-1* promoter, and the *rab-3* promoter was inserted using the In-Fusion HD Cloning Kit (Takara Bio). The resultant *rab-3p::frpr-9 DNA* plasmid was injected at 10ng/µl, with 50ng/µl pCFJ2474 (Cas9 expression), 50ng/µl pJW1882 (ttTi4348 targeting sequence) and 5ng/µl *myo-2p::mCherry into frpr-9(sy1294)* animals.

*gcy-9p::gcy-9* single copy rescue construct

The 3246bp *gcy-9* cDNA was amplified from a *C. elegans* cDNA library with forward (TGAGATGATAAATCGATATGCGTTTATATTTATTTTTC) and reverse (GCATTTATCTTAGCCGGCTCATTGTTTGCCGGTTCTTCC) primers from Sigma-Aldrich incorporating *Bsu15I-PdiI* restriction sites. The *gcy-9p-FLP-p2A-H2B-mTurq2* plasmid was digested using *Bsu15I-PdiI* and the *gcy-9* cDNA was inserted using the In-Fusion HD Cloning Kit (Takara Bio). The resultant *gcy-9p::gcy-9 cDNA* plasmid was injected at 10ng/µl, with 50ng/µl pCFJ2474 (Cas9 expression), 50ng/µl pJW1882 (ttTi4348 targeting sequence) and 5ng/µl *myo-2p::mCherry into gcy-9(n4470); hsp-6p::gfp* and *gcy-9(n4470); hsp-60p::gfp* animals.

**Auxin inducible degradation**

35mm NGM plates were supplemented with 1 mM auxin (indole-3-acetic acid, catalog ALFA10556.14, Thermo Scientific) from a stock of 400mM dissolved in ethanol. NGM plates supplemented with ethanol were used as a control. Plates were seeded with 400µl OP50. 5 L4 animals were placed on each plate and after 5 days L4 progeny of the next generation were analysed.

**Acute paraquat sensitivity assays**

35mm NGM plates were supplemented with 200mM paraquat (methyl viologen dichloride hydrate, catalog 856177, Sigma Aldrich) *(43)*. Plates were dried overnight at room temperature, then seeded with 50µl 10x concentrated OP50. Bacteria was grown at room temperature overnight. 25 L4 worms were then transferred to each plate with an eyebrow hair, and the survival of the animals was assessed each hour for 15 consecutive hours. Worms that left the agar were excluded from the analysis.

**TMRE staining**

TMRE (Tetramethylrhodamine Ethyl Ester Perchlorate) staining was adapted from previously described methods *(43,44)*. In short, 100µl of M9 containing 1mM TMRE was seeded over the top of the 400µl OP50 on NGM plates. Plates were covered and dried at room temperature. L2 hermaphrodites were placed on the plates and left to stain for 20 hours. Worms were removed from TMRE plates and de-stained on NGM plates seeded with 400µl OP50 for four hours to clear excess dye from the intestinal cavity. Live L4 animals were then imaged, and relative fluorescence was calculated as described above.

**Oxygen consumption rate assays**

Oxygen consumption rate was measured using the Seahorse XFe96 extracellular flux analyser, as described previously *(46)*. The sensor cartridge was hydrated with Seahorse XF Calibrant solution overnight in a $CO_2$ free 37°C incubator. Injection ports were loaded with 25µl 90µM FCCP (port A) and 25µl 500mM sodium azide (port B) for every well. 100 L4 animals of each genotype were picked to empty NGM plates, washed in M9, and divided between 6 wells of a cell culture plate for technical replicates, at a final volume of 200µl per well. The exact number of worms per well was counted for normalisation. Wells without worms were filled with 200µl M9. The sensor cartridge was calibrated, and then mitochondrial respiration was measured using the following protocol:
Basal respiration measurement, loop 6 times:

- Mix for 3 minutes
- Hold for 2 minutes
- Measure for 3 minutes

Maximal respiration measurement, inject port A (FCCP), loop 6 times:

- Mix for 3 minutes
- Hold for 2 minutes
- Measure for 3 minutes

Non-mitochondrial respiration measurement, inject port B (sodium azide), loop 6 times:

- Mix for 3 minutes
- Hold for 2 minutes
- Measure for 3 minutes
- Loop end

Respiration was normalised to the number of worms per well and averaged across the technical replicates. The experiment was performed twice over two separate days to confirm reproducibility.

**Oil-red O (ORO) fat staining**

ORO fat staining was performed as described previously *(18)*. Briefly, ORO stock solution (0.5g ORO dissolved in 100ml isopropanol) was diluted to 60% in milli-Q water. ~600 synchronised day 1 adult worms were washed 3 times with 1XPBS, then resuspended in 1000µl 60% isopropanol and fixed at room temperature for 20 minutes. Samples were centrifuged at 3300 RCF for 30 seconds and the supernatant was removed. The 60% Oil-Red O solution was then filtered through a 0.22µm mesh filter. 400µl of the filtered solution was added to each tube containing a worm pellet, which were covered in foil and incubated at room temperature overnight on a rotating rack.
The following morning, stained worms were washed twice with 500µl PBS + 0.01% Triton-X100 and once with 1XPBS. After the final wash step, worms were resuspended in a small amount of remaining 1XPBS and pipetted onto a 5% agarose pad on a glass slide for imaging. Pseudo-RGB images were taken of the four most proximal intestinal cells using the 40x objective. The pharynx was used as the focal point for images to ensure consistent focal planes between images. Staining intensity was measured using ImageJ software by tracing the first four intestinal cells. As Oil-Red O absorbs green light, the staining intensity is measured using

the inverted green channel. A region outside of the worm was used as a background measurement. CTCF was calculated as described above.

**Food deprivation assay**

OP50 E. coli was cultured overnight to saturation. To generate dead bacteria, the overnight OP50 culture was treated with 4% paraformaldehyde (PFA) solution to a final concentration of 0.5% PFA, and incubated at 37ºC for one hour *(47)*. Bacterial culture was washed three times in LB to remove residual PFA. The bacterial pellet was resuspended in one fifth of the original volume to generate a 5x dilution. 500ul of either live or dead OP50 was uniformly seeded on NGM agar plates and allowed to dry overnight. L4 animals were transferred to uniformly seeded OP50 (live bacteria), PFA-treated OP50 (dead bacteria) or empty plates (no bacteria) and incubated at 20ºC for 24 hours prior to imaging.

**Statistics and reproducibility**

Experiments were performed in three independent replicates, unless otherwise stated in the text. The experimenter was blinded to genotype/condition. Statistical tests were performed in GraphPad Prism 10 software, using either a Welch's t test when comparing two samples, one-way ANOVA for comparing more than two samples or two-way ANOVA for survival assays, as indicated in figure legends. Values are expressed as mean ± SEM. Differences with a $p$ value <0.05 were considered significant. The number of animals analysed and exact $p$ values are reported in each figure legend.

### Supplementary Figures

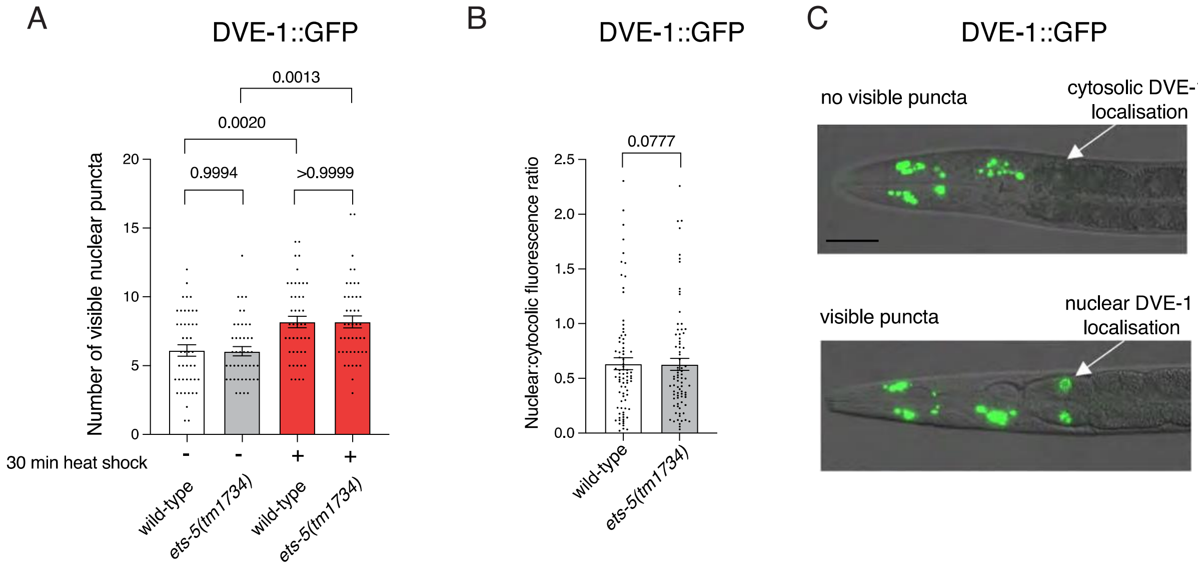


**Fig. S1. DVE-1 is not activated by *ets-5* loss.**

(**A**) Quantification of DVE-1::GFP nuclear puncta in wild-type and *ets-5(tm1734)* animals with and without 30 minutes heat shock. *n* = 45. (**B**) Quantification of DVE-1::GFP nuclear:cytosolic ratio in wild-type and *ets-5(tm1734)* animals. *n* = 60. (**F**) DIC and fluorescent micrographs of DVE-1::GFP, showing no visible (top) and visible (bottom) nuclear puncta in the first intestinal cells. *P* values assessed by one-way analysis of variance (ANOVA) with Tukey's post hoc test (**A**), or unpaired t test with Welch's correction (**B**). Error bars indicate SEM. Scale bar, 50μm.

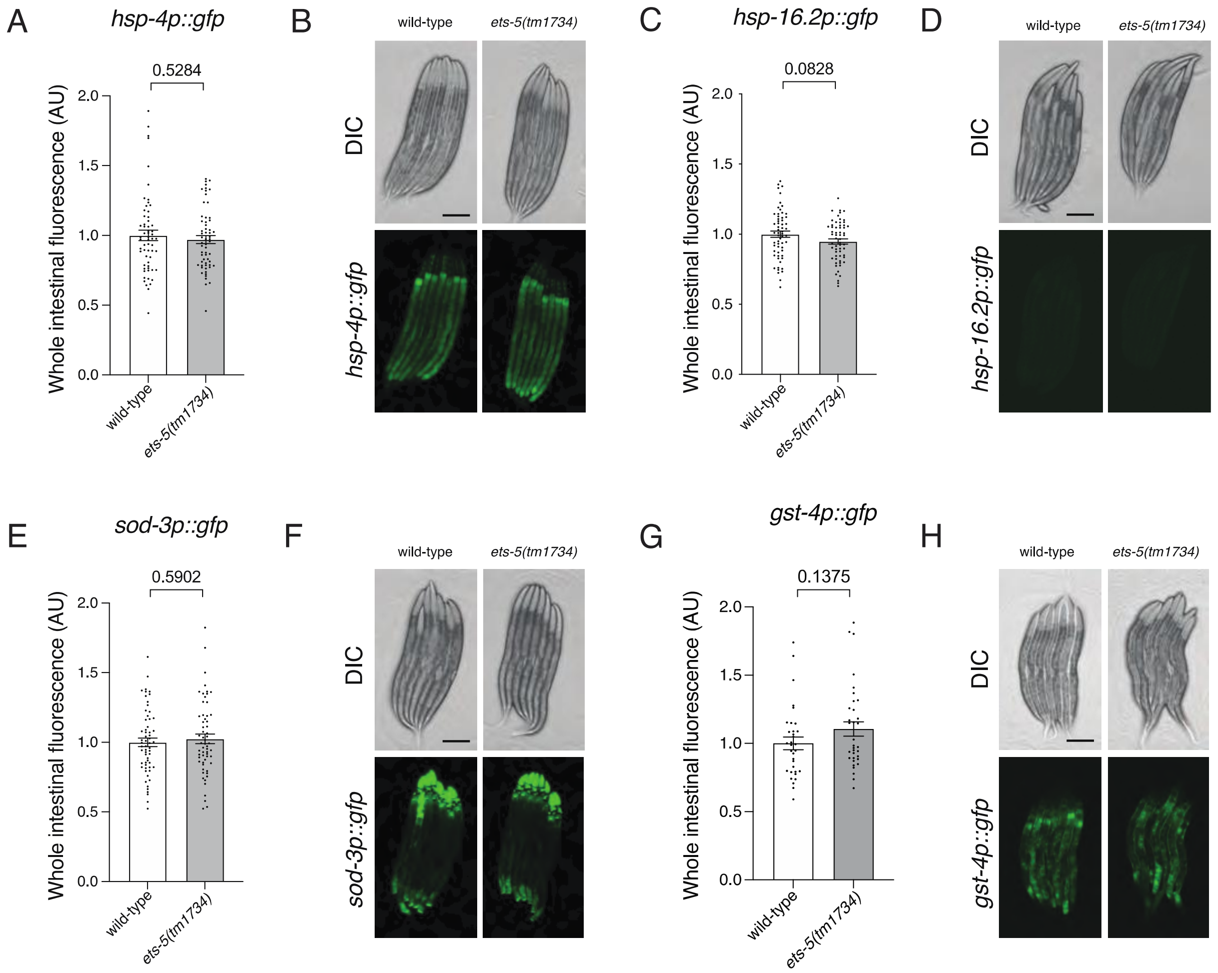


**Fig. S2. ETS-5 intestinal stress response regulation is limited to the UPR$^{mt}$.**

**A, B** Quantification (**A**) and DIC/fluorescent micrographs (**B**) of UPR$^{ER}$ reporter (*hsp-4p::gfp*) expression in L4 larvae of wild-type and *ets-5(tm1734)* animals. *n* = 60 **C, D** Quantification (**C**) and DIC/fluorescent micrographs (**D**) of cytosolic heat shock response reporter (*hsp-16.2p::gfp*) expression in L4 larvae of wild-type and *ets-5(tm1734)* animals. *n* = 70 **E, F** Quantification (**E**) and DIC/fluorescent micrographs (**F**) of oxidative stress response reporter (*sod-3p::gfp*) expression in L4 larvae of wild-type and *ets-5(tm1734)* animals. *n* = 60 **G, H** Quantification (**G**) and DIC/fluorescent micrographs (**H**) of oxidative stress response reporter (*gst-4p::gfp*) expression in L4 larvae of wild-type and *ets-5(tm1734)* animals. *n* = 30. *P* values assessed by unpaired t test with Welch's correction. Error bars indicate SEM. Scale bars, 250µm.

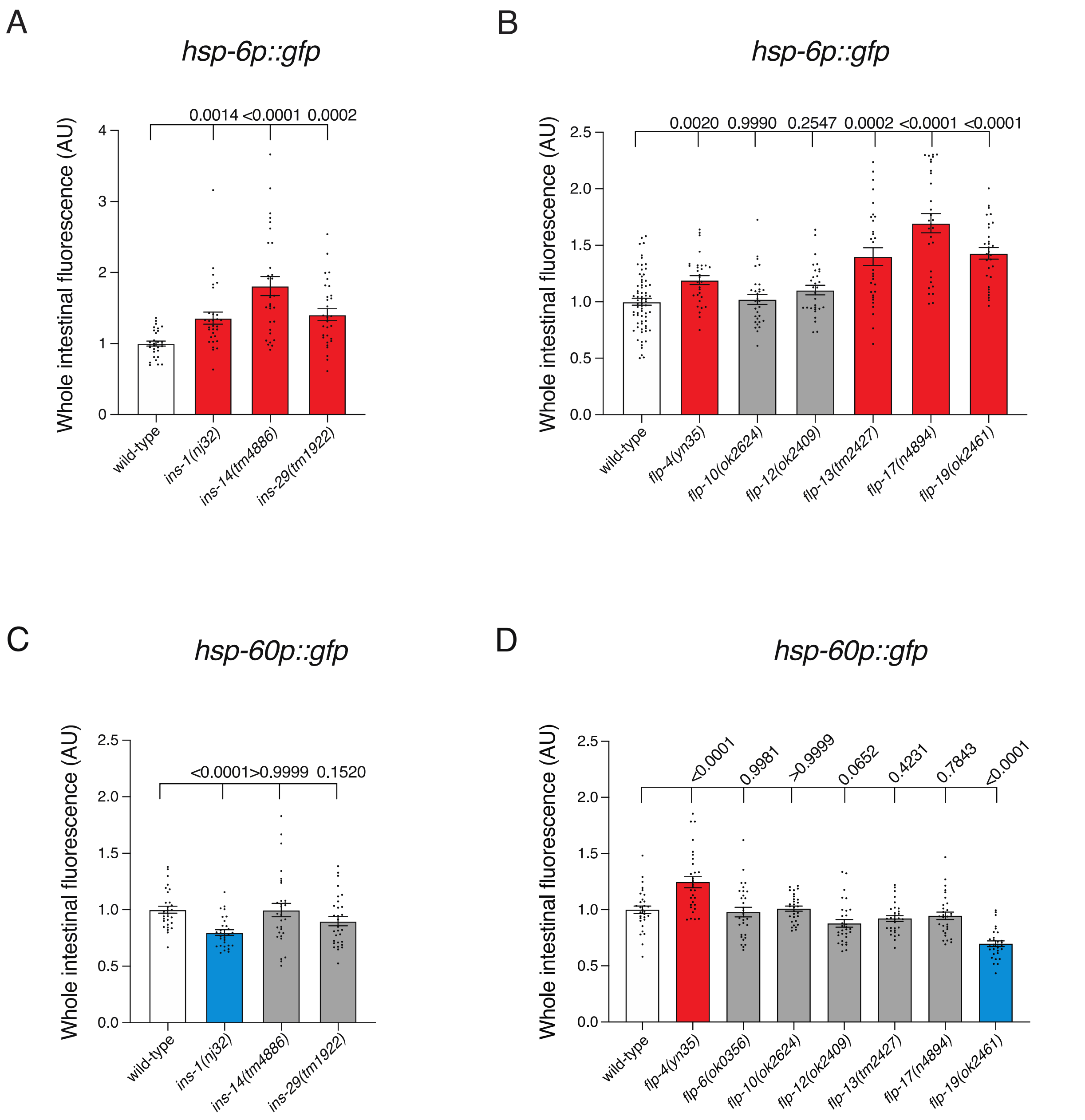


**Fig. S3. Neuropeptides expressed by the BAG neurons differentially impact intestinal HSP-6 and HSP-60 expression.**

**A, B** Quantification of UPR$^{mt}$ reporter (*hsp-6p::gfp*) expression in L4 larvae of wild-type and (**A**) *ins-1(nj32), ins-14(tm4886)* and *ins-29(tm1922)* animals, and (**B**) *flp-4(yn35), flp-10(ok2624), flp-12(ok2409), flp-13(tm2427), flp-17(n4894)* and *flp-19(ok2461)* animals. **C, D** Quantification of UPR$^{mt}$ reporter (*hsp-60p::gfp*) expression in L4 larvae of wild-type and (**C**) *ins-1(nj32), ins-14(tm4886)* and *ins-29(tm1922)* animals, and (**D**) *flp-4(yn35), flp-6(3056), flp-10(ok2624), flp-12(ok2409), flp-13(tm2427), flp-17(n4894)* and *flp-19(ok2461)* animals. $n$ = 30. $P$ values assessed by one-way analysis of variance (ANOVA) with Tukey's post hoc test. Error bars indicate SEM.

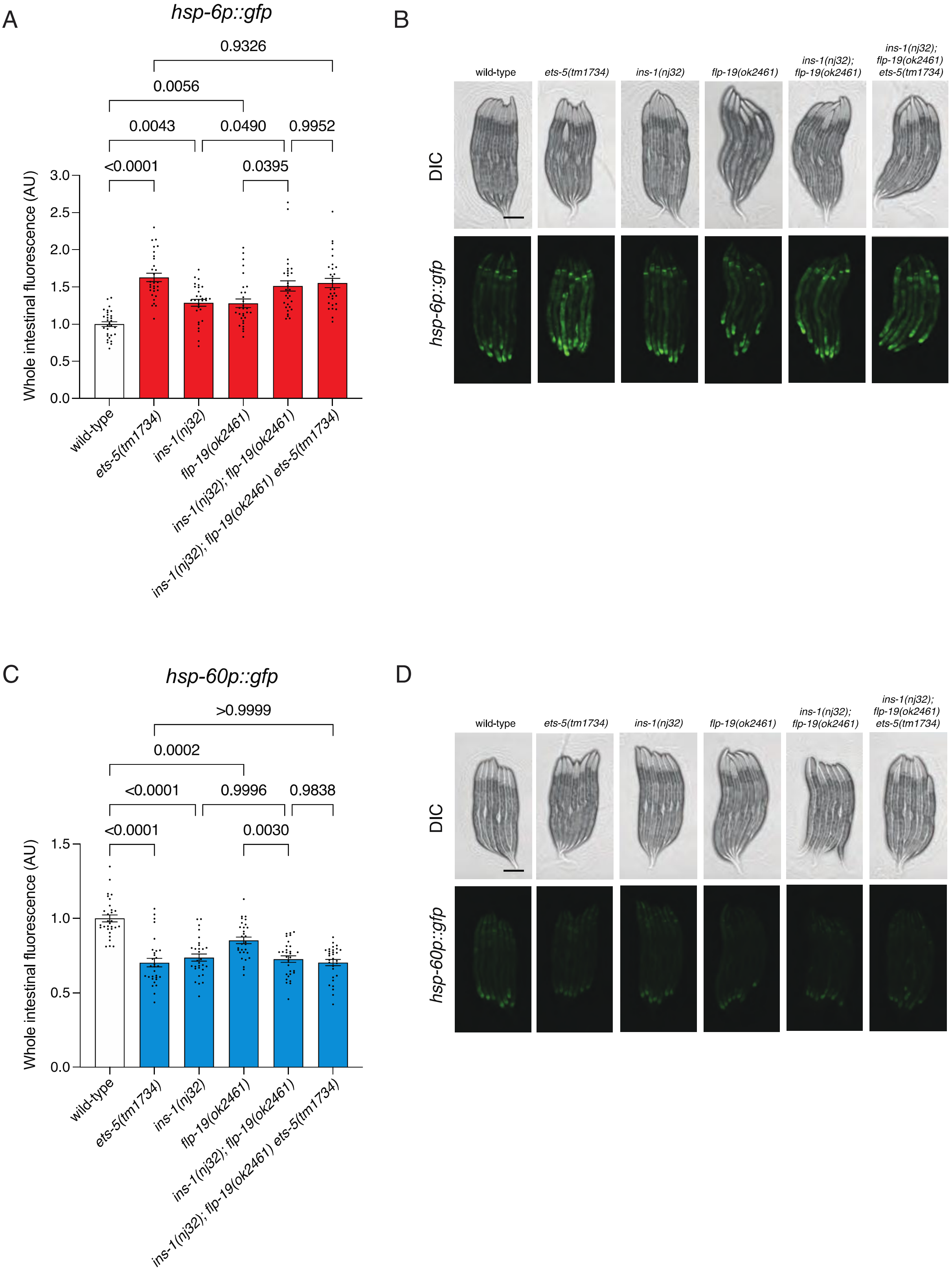
A
hsp-6p::gfp
0.9326
0.0056
0.0043
0.0490
0.9952
<0.0001
0.0395
Whole intestinal fluorescence (AU)
3.0
2.5
2.0
1.5
1.0
0.5
0.0
wild-type
ets-5(tm1734)
ins-1(nj32)
flp-19(ok2461)
ins-1(nj32); flp-19(ok2461)
ins-1(nj32); flp-19(ok2461) ets-5(tm1734)
B
wild-type
ets-5(tm1734)
ins-1(nj32)
flp-19(ok2461)
ins-1(nj32); flp-19(ok2461)
ins-1(nj32); flp-19(ok2461) ets-5(tm1734)
DIC
hsp-6p::gfp
C
hsp-60p::gfp
>0.9999
0.0002
<0.0001
0.9996
0.9838
<0.0001
0.0030
Whole intestinal fluorescence (AU)
1.5
1.0
0.5
0.0
wild-type
ets-5(tm1734)
ins-1(nj32)
flp-19(ok2461)
ins-1(nj32); flp-19(ok2461)
ins-1(nj32); flp-19(ok2461) ets-5(tm1734)
D
wild-type
ets-5(tm1734)
ins-1(nj32)
flp-19(ok2461)
ins-1(nj32); flp-19(ok2461)
ins-1(nj32); flp-19(ok2461) ets-5(tm1734)
DIC
hsp-60p::gfp

**Fig. S4. ETS-5 mediates a non-canonical systemic UPR$^{mt}$ in the same pathway as neuropeptides INS-1 and FLP-19.**

**A, B** Quantification (**A**) and DIC/fluorescent micrographs (**B**) of UPR$^{mt}$ reporter (*hsp-6p::gfp*) expression in L4 larvae of wild-type, *ets-5(tm1734), ins-1(nj32), flp-19(ok2461)*, *ins-1(nj32); flp-19(ok2461)* and *ins-1(nj32); flp-19(ok2461) ets-5(tm1734)* animals. **C, D** Quantification (**C**) and DIC/fluorescent micrographs (**D**) of UPR$^{mt}$ reporter (*hsp-60p::gfp*) expression in L4 larvae of wild-type, *ets-5(tm1734), ins-1(nj32), flp-19(ok2461)*, *ins-1(nj32); flp-19(ok2461)* and *ins-1(nj32); flp-19(ok2461) ets-5(tm1734)* animals. $n$ = 30.$P$ values assessed by one-way analysis of variance (ANOVA) with Tukey's post hoc test. Error bars indicate SEM. Scale bars, 250μm.

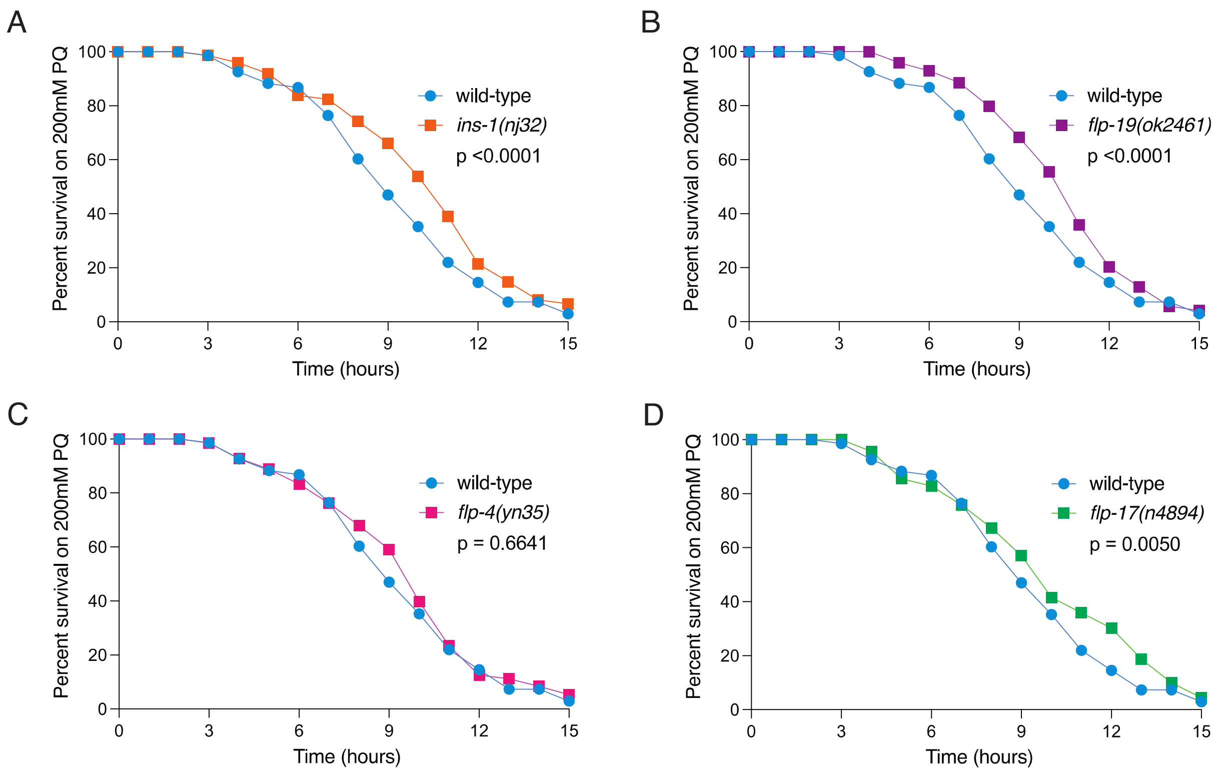


**Fig. S5. INS-1 and FLP-19 mutants are resistant to mitochondrial stress.**

Survival analysis of wild-type (*n*=67), (**A**) *ins-1(nj32)* (*n*=74), (**B**) *flp-19(ok2461)* (*n*=70), (**C**) *flp-4(yn35)* (*n*=72) and (**D**) *flp-17(n4894)* (*n*=70) animals exposed to 200mM paraquat from the L4 larval stage. *P* values assessed by two-way analysis of variance (ANOVA) with Tukey's post hoc test.

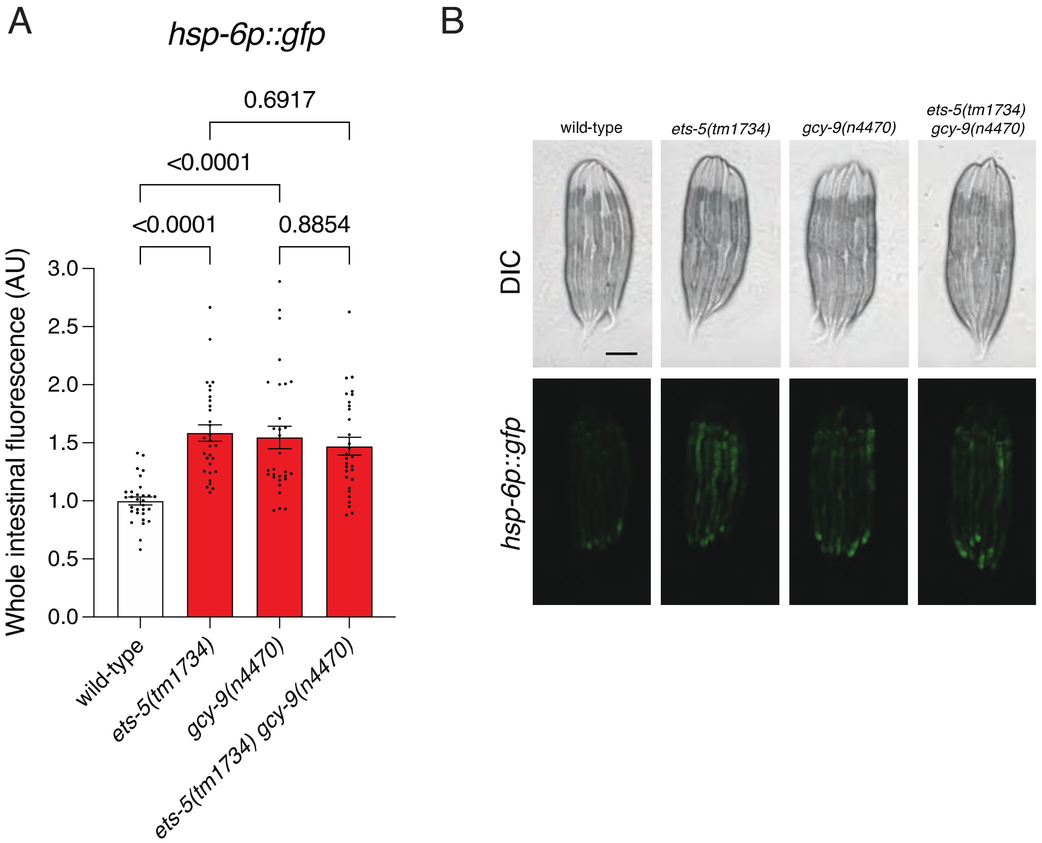
A
hsp-6p::gfp
0.6917
<0.0001
<0.0001
0.8854
Whole intestinal fluorescence (AU)
3.0
2.5
2.0
1.5
1.0
0.5
0.0
wild-type
ets-5(tm1734)
gcy-9(n4470)
ets-5(tm1734) gcy-9(n4470)
B
wild-type
ets-5(tm1734)
gcy-9(n4470)
ets-5(tm1734)
gcy-9(n4470)
DIC
hsp-6p::gfp

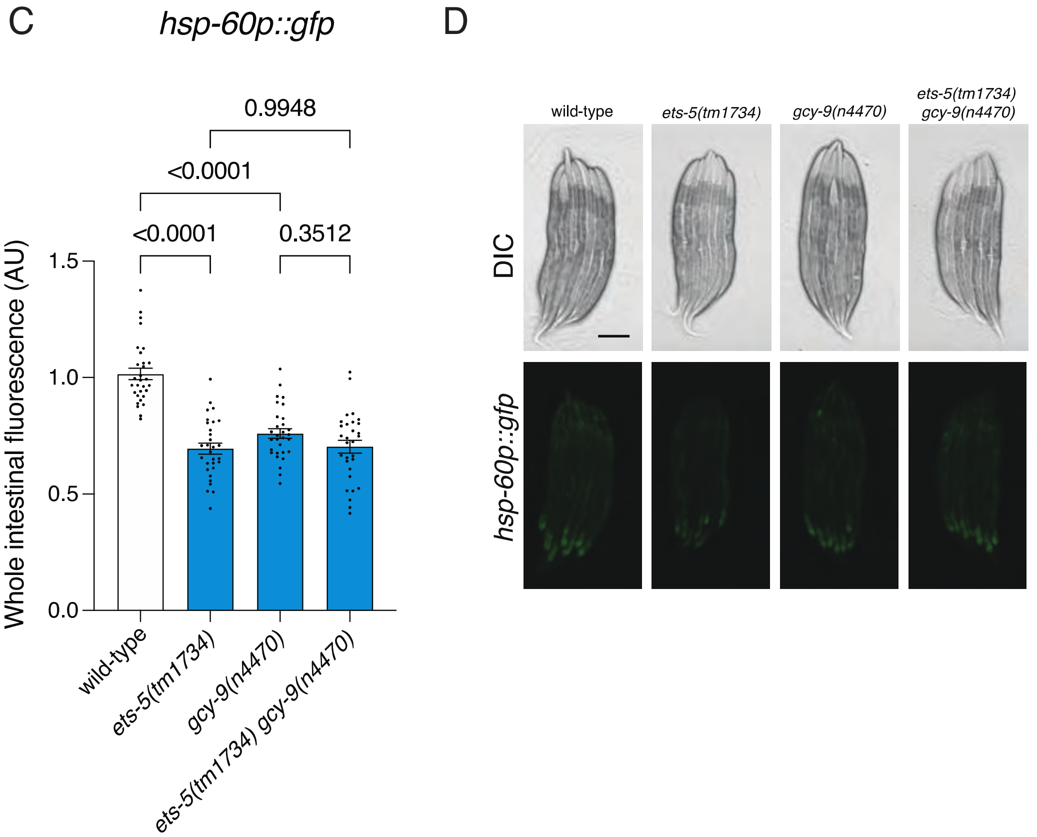
C
hsp-60p::gfp
0.9948
<0.0001
<0.0001
0.3512
Whole intestinal fluorescence (AU)
1.5
1.0
0.5
0.0
wild-type
ets-5(tm1734)
gcy-9(n4470)
ets-5(tm1734) gcy-9(n4470)
D
wild-type
ets-5(tm1734)
gcy-9(n4470)
ets-5(tm1734)
gcy-9(n4470)
DIC
hsp-60p::gfp

**Fig. S6. ETS-5 and GCY-9 function in the same pathway to mediate a non-canonical systemic UPR$^{mt}$.**

**A, B** Quantification (**A**) and DIC/fluorescent micrographs (**B**) of UPR$^{mt}$ reporter (*hsp-6p::gfp*) expression in L4 larvae of wild-type, *ets-5(tm1734)*, *gcy-9(n4470)* and *ets-5(tm1734) gcy-9(n4470)* animals. **C, D** Quantification (**C**) and DIC/fluorescent micrographs (**D**) of UPR$^{mt}$ reporter (*hsp-60p::gfp*) expression in L4 larvae of wild-type, *ets-5(tm1734)*, *gcy-9(n4470)* and *ets-5(tm1734) gcy-9(n4470)* animals. $n$ = 30. $P$ values assessed by one-way analysis of variance (ANOVA) with Tukey's post hoc test. Error bars indicate SEM. Scale bars, 250μm.

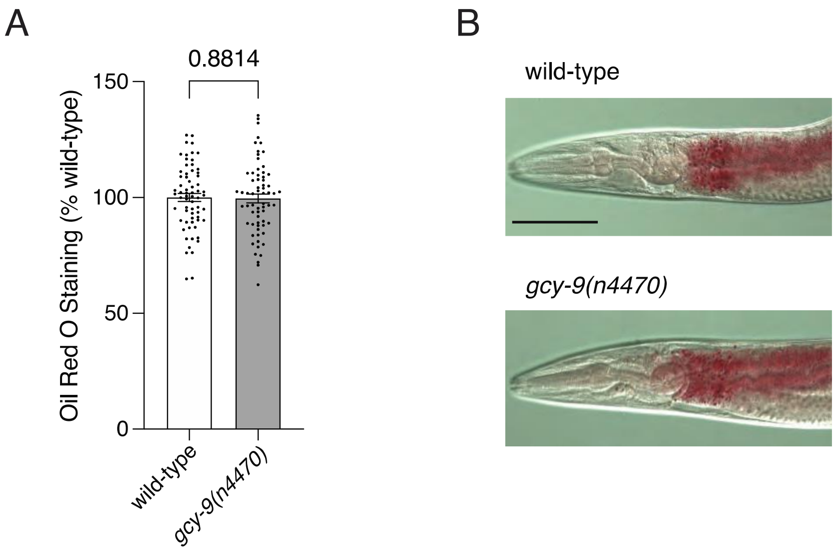


**Fig. S7. Loss of *gcy-9* does not impact intestinal fat levels.**

**A, B** Quantification (**A**) and representative images (**B**) of Oil Red O staining in wild-type and *gcy-9(n4470)* adult hermaphrodites. *P* value assessed or unpaired t test with Welch's correction. Error bars indicate SEM. *n* = 60. Scale bars, 50μm.

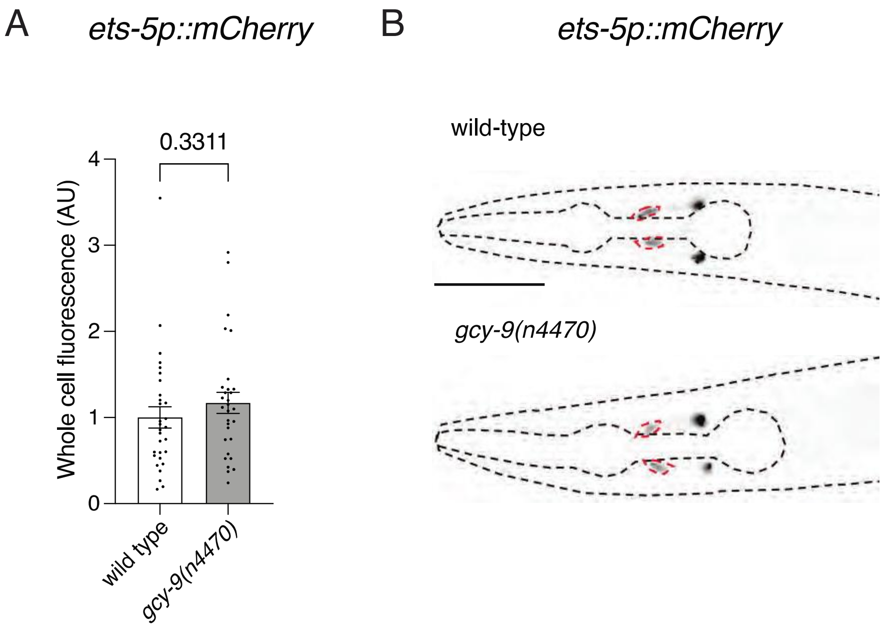


**Fig. S8. Loss of *gcy-9* does not impact *ets-5* expression.**

**A, B** Quantification (**A**) and fluorescent micrographs (**B**) of *ets-5p::mCherry* reporter expression in the BAG neurons in L4 larvae of wild-type and *gcy-9(n4470)* animals. *n* = 30. Scale bar, 50µm.

**Table S1. *C. elegans* strains used in this work.**

| Strain name | Genotype | Source |
|---|---|---|
| SJ4100 | *zcIs13[hsp-6p::GFP + lin-15(+)]V* | CGC |
| RJP5477 | *zcIs13[hsp-6p::GFP + lin-15(+)]V; ets-5(tm1734)X* | this study |
| RJP5337 | *zcIs13[hsp-6p::GFP + lin-15(+)]V; ets-5(tm1755)X* | this study |
| RJP5838 | *zcIs13[hsp-6p::GFP + lin-15(+)]V; ets-5(tm1734)X; rpEx2159[gcy-9p::ets-5 cDNA]* | this study |
| SJ4058 | *zcIs9[hsp-60p::GFP + lin-5(+)]V* | CGC |
| RJP4183 | *zcIs9[hsp-60p::GFP + lin-5(+)]V; ets-5(tm1734)X* | this study |
| RJP5791 | *zcIs9[hsp-60p::GFP + lin-5(+)]V; ets-5(tm1755)X* | this study |
| RJP5931 | *zcIs9[hsp-60p::GFP + lin-5(+)]V; ets-5(tm1734)X; rpEx2159[gcy-9p::ets-5 cDNA]* | this study |
| RJP4821 | *unc-25(e156)III; zcIs13[hsp-6p::GFP + lin-15(+)]V* | Pocock lab |
| SJ4197 | *zcIs31[dve-1p::dve-1::GFP]II* | CGC |
| RJP4398 | *zcIs31[dve-1p::dve-1::GFP]II; ets-5(tm1734)X* | this study |
| Bristol strain N2 | wild type | CGC |
| RJP235 | *ets-5(tm1734)X* | NBRP |
| RJP5930 | *ets-5(tm1734)X; rpEx2159[gcy-9p::ets-5 cDNA]* | this study |
| RJP5590 | *rpSi1[gcy-9p::TIR1::F2A::mTagBFP2::NLS::AID::tbb-2 3'UTR]II; rp166[unc-31-linker-GFP-TEV-AID-FLAG]IV; zcIs13[hsp-6p::GFP + lin-15(+)]V* | this study |
| RJP5654 | *rpSi1[gcy-9p::TIR1::F2A::mTagBFP2::NLS::AID::tbb-2 3'UTR]II; rp166[unc-31-linker-GFP-TEV-AID-FLAG]IV; zcIs9[hsp-60p::GFP + lin-5(+)]V* | this study |
| RJP4435 | *ins-1(nj32)IV; zcIs13[hsp-6p::GFP + lin-15(+)]V* | this study |
| RJP7040 | *rp306[gcy-9p::ins-1 genomic DNA]I; ins-1(nj32)IV; zcIs13[hsp-6p::GFP + lin-15(+)]V* | this study |
| RJP5028 | *zcIs13[hsp-6p::GFP + lin-15(+)]V; flp-19(ok2461)X* | this study |
| RJP7042 | *rp308[gcy-9p::flp-19 genomic DNA]I; zcIs13[hsp-6p::GFP + lin-15(+)]V; flp-19(ok2461)X* | this study |
| RJP4980 | *ins-1(nj32)IV; zcIs9[hsp-60p::GFP + lin-5(+)]V* | this study |
| RJP7041 | *rp307[gcy-9p::ins-1 genomic DNA]I; ins-1(nj32)IV; zcIs9[hsp-60p::GFP + lin-5(+)]V* | this study |
| RJP6089 | *zcIs9[hsp-60p::GFP + lin-5(+)]V; flp-19(ok2461)X* | this study |
| RJP7044 | *rp309[gcy-9p::flp-19 genomic DNA]I; zcIs9[hsp-60p::GFP + lin-5(+)]V; flp-19(ok2461)X* | this study |
| IK581 | *ins-1(nj32)IV* | CGC |
| RB1903 | *flp-19(ok2461)X* | CGC |
| RJP7020 | *ins-1(nj32)IV; flp-19(ok2461)X* | this study |
| RJP7021 | *ins-1(nj32)II; flp-19(ok2461) ets-5(tm1734)X* | this study |
| PS8398 | *frpr-9(sy1294)V* | CGC |
| RJP7394 | *frpr-9(sy1294)V; flp-19(n2461)X* | this study |

| | | |
|---|---|---|
| RJP7170 | *rp356(ges-1p::frpr-9 genomic DNA)I; frpr-9(sy1294)V* | this study |
| RJP7225 | *rp366(rab-3p::frpr-9 genomic DNA)I; frpr-9(sy1294)V* | this study |
| RJP4102 | *rpEx1752[ins-1p::gfp + rol-6]* | Pocock lab |
| RJP7050 | *gcy-9(n4470)X; rpEx1752[ins-1p::gfp + rol-6]* | this study |
| RJP7046 | *gcy-9(n4470)X* | CGC |
| RJP7047 | *ets-5(tm1734) gcy-9(n4470)X* | this study |
| RJP7048 | *zcIs13[hsp-6p::GFP + lin-15(+)]V; gcy-9(n4470)X* | this study |
| RJP7396 | *rp406{gcy-9p::gcy-9]I; zcIs13[hsp-6p::GFP + lin-15(+)]V; gcy-9(n4470)X* | this study |
| RJP7049 | *zcIs9[hsp-60p::GFP + lin-5(+)]V; gcy-9(n4470)X* | this study |
| RJP7328 | *rp398[gcy-9p::gcy-9]I; zcIs9[hsp-60p::GFP + lin-5(+)]V; gcy-9(n4470)X* | this study |
| FQ383 | *gcy-9p::gfp* | Ringstad lab |
| SJ4005 | *zcIs4[hsp-4p::gfp]V* | CGC |
| RJP4171 | *zcIs4[hsp-4p::gfp]V; ets-5(tm1734)X* | this study |
| CL2070 | *dvIs70[hsp-16.2p::GFP + rol-6(su1006)]* | CGC |
| RJP4166 | *ets-5(tm1734)X; dvIs70[hsp-16.2p::GFP + rol-6(su1006)]* | this study |
| CF1553 | *muIs84[sod-3p::GFP + rol-6(su1006)]* | CGC |
| RJP4399 | *ets-5(tm1734)X; muIs84[sod-3p::GFP + rol-6(su1006)]* | this study |
| CL2166 | *dvIs19[(pAF15)gst-4p::gfp::nls]III* | CGC |
| RJP7034 | *dvIs19[(pAF15)gst-4p::gfp::nls]III; ets-5(tm1734)X* | this study |
| RJP4971 | *ins-14(tm4886)II; (nj32)IV; zcIs13[hsp-6p::GFP + lin-15(+)]V* | this study |
| RJP4976 | *ins-29(tm1922)I; (nj32)IV; zcIs13[hsp-6p::GFP + lin-15(+)]V* | this study |
| RJP5107 | *flp-4(yn35)II; (nj32)IV; zcIs13[hsp-6p::GFP + lin-15(+)]V* | this study |
| RJP5060 | *flp-10(ok2624)IV; (nj32)IV; zcIs13[hsp-6p::GFP + lin-15(+)]V* | this study |
| RJP4978 | *zcIs13[hsp-6p::GFP + lin-15(+)]V; flp-12(2409)X* | this study |
| RJP4986 | *flp-13(tm2427)IV; (nj32)IV; zcIs13[hsp-6p::GFP + lin-15(+)]V* | this study |
| RJP5166 | *flp-17(n4894)IV; (nj32)IV; zcIs13[hsp-6p::GFP + lin-15(+)]V* | this study |
| RJP5034 | *ins-14(tm4886)II; zcIs9[hsp-60p::GFP + lin-5(+)]V* | this study |
| RJP4973 | *ins-29(tm1922)I; zcIs9[hsp-60p::GFP + lin-5(+)]V* | this study |
| RJP5108 | *flp-4(yn35)II; zcIs9[hsp-60p::GFP + lin-5(+)]V* | this study |
| RJP5030 | *flp-6(ok3056)V; zcIs9[hsp-60p::GFP + lin-5(+)]V* | this study |
| RJP5017 | *flp-10(ok2624)IV; zcIs9[hsp-60p::GFP + lin-5(+)]V* | this study |
| RJP4973 | *zcIs9[hsp-60p::GFP + lin-5(+)]V; flp-12(2409)X* | this study |
| RJP5051 | *flp-13(tm2427)IV; zcIs9[hsp-60p::GFP + lin-5(+)]V* | this study |

| | | |
|---|---|---|
| RJP5109 | *flp-17(n4894)IV; zcIs9[hsp-60p::GFP + lin-5(+)]V* | this study |
| RJP6201 | *ins-1(nj32)IV; zcIs13[hsp-6p::GFP + lin-15(+)]V; flp-19(ok2461)X* | this study |
| RJP7022 | *ins-1(nj32)IV; zcIs13[hsp-6p::GFP + lin-15(+)]V; flp-19(ok2461) ets-5(tm1734)X* | this study |
| RJP7026 | *ins-1(nj32)IV; zcIs9[hsp-60p::GFP + lin-5(+)]V; flp-19(ok2461)X* | this study |
| RJP7027 | *ins-1(nj32)IV; zcIs9[hsp-60p::GFP + lin-5(+)]V; flp-19(ok2461; ets-5(tm1734))X* | this study |
| PS9050 | *flp-4(yn35)II* | CGC |
| MT15933 | *flp-17(n4894)IV* | CGC |
| RJP7171 | *zcIs13[hsp-6p::GFP + lin-15(+)]V; gcy-9(n4470) ets-5(tm1734)X* | this study |
| RJP7172 | *zcIs9[hsp-60p::GFP + lin-5(+)]V; ] gcy-9(n4470) ets-5(tm1734)X* | this study |
| RJP3085 | *rpEx1520[ets-5p::mCherry]* | Pocock lab |
| RJP7181 | *gcy-9(n4470)X; rpEx1520[ets-5p::mCherry]* | this study |

**Data S1. (separate file)**

Source Data.
